# 3D imaging of human pancreas suggests islet size and endocrine composition influence their loss in type 1 diabetes

**DOI:** 10.1101/2025.05.14.654045

**Authors:** Alexandra Rippa, Amanda L. Posgai, Seth Currlin, Maigan Brusko, MacKenzie D. Williams, John S. Kaddis, Irina Kusmartseva, Clive H. Wasserfall, Martha Campbell-Thompson, Mark A. Atkinson

## Abstract

A high-definition description of pancreatic islets would prove beneficial for understanding the pathophysiology of type 1 diabetes (T1D), yet significant knowledge voids exist in terms of their size, endocrine cell composition, and number in both health and disease. Here, 3-dimensional (3D) analyses of pancreata from control persons without diabetes (ND) revealed heretofore underappreciated frequencies (approximately 50%) of insulin-positive (INS+) glucagon-negative (GCG-) islets. Non-diabetic individuals positive for a single Glutamic acid decarboxylase autoantibody (GADA+) yet at increased risk for disease consistently demonstrated endocrine features, including islet volume and cell composition, closely resembling the age-matched ND controls. In contrast, pancreata from individuals with short-duration T1D demonstrated significantly reduced islet density and a dramatic loss of INS+GCG- islets with preservation of large INS+GCG+ islets. The size and cellular composition of pancreatic islets may, therefore, represent influential factors that impact β-cell loss during T1D disease progression.

**Graphical Abstract:** 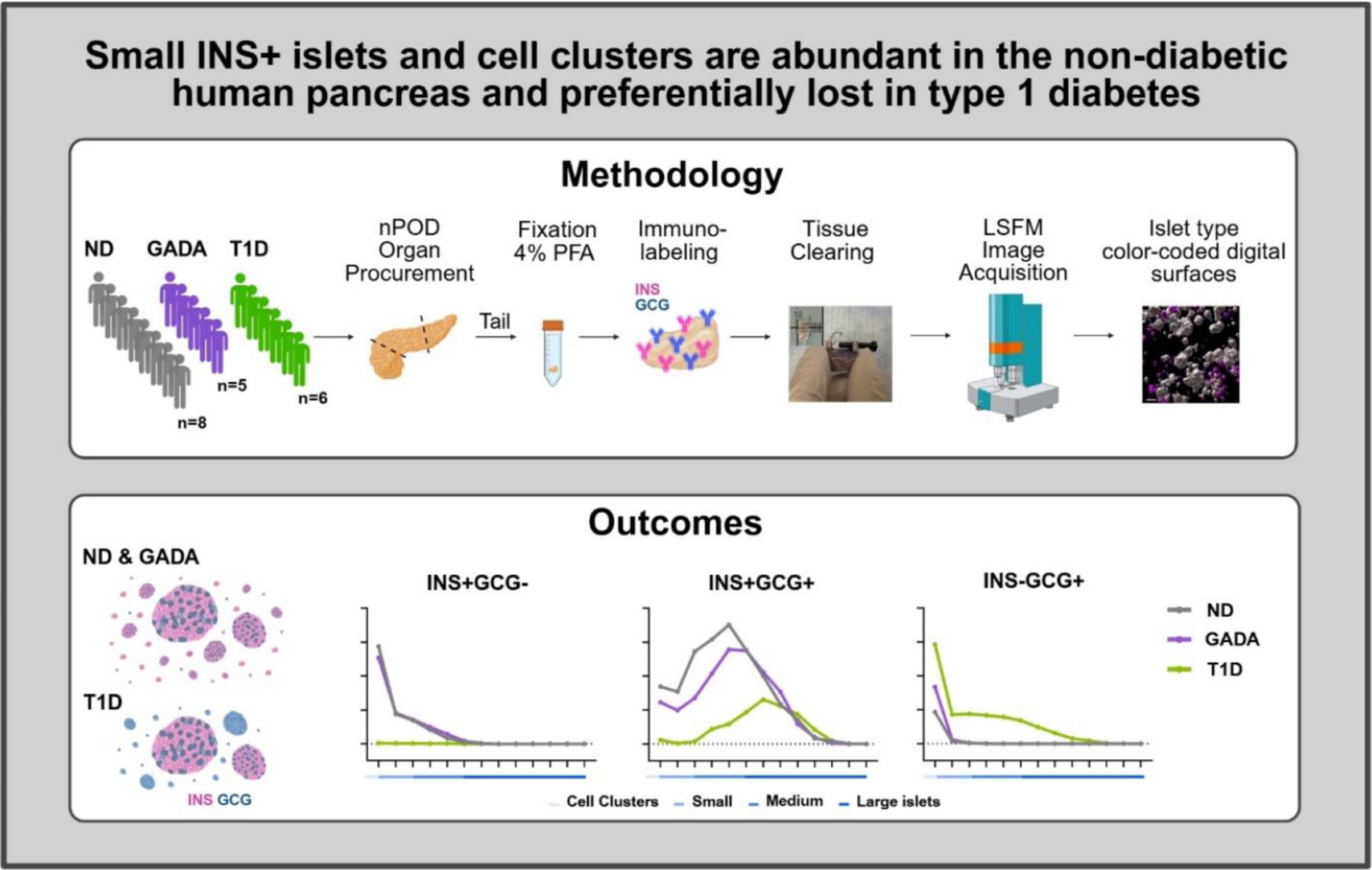

**Highlights:**

- 3D imaging of non-diabetic (ND) pancreas suggests up to 50% of INS+ islets lack GCG
- INS+GCG- islets are typically small-sized in ND and preferentially lost in T1D
- In T1D pancreas, INS+ beta cells are preserved in large INS+GCG+ islets
- Islets from GADA+ individuals appear similar to ND for size and β-to-α cell ratios

## INTRODUCTION

The pancreatic islets of Langerhans, which play a crucial role in glucose regulation through endocrine hormone secretion, have long been a focus of metabolic research due to their impact on health and disease, particularly in diabetes^1^. These endocrine micro-organs are surrounded by a complex network of blood and lymphatic vessels, ducts, neurons, and extracellular matrix; however, estimates of pancreatic islet quantity, volume, and cellular composition vary widely in the literature^2–4^. While some of this variation can be attributed to individual anthropometric factors, such as age and body mass index^5^, differences in analytical methods—ranging from two-dimensional (2D) microscopy^2–4^ to positron emission tomography (PET) imaging *in vivo*^6,7^, studies of isolated islets^8^, live pancreas slices^9^, and more—have also contributed to quantitative data inconsistencies.

Commonly cited estimates of islet quantity as a percentage of total pancreatic mass routinely range from 1 to 2%, with absolute numbers varying dramatically between 3.2 and 14.8 million islets per pancreas^10^. Similarly, estimates of islet size show considerable variability, ranging from approximately 30 to >400 µm in diameter (with reported mean diameters varying from 65-110 μm across 2D and 3-dimensional (3D) imaging modalities)^11–13^ and from 2.5x10^4^ to 5.1x10^7^ µm^3^ in volume^13,14^. However, until recently, endocrine cell clusters <30 µm in diameter were often excluded from qualitative or quantitative analyses. For decades, islet endocrine cell composition, determined with 2D based methods, was reported as approximating 60% insulin (INS)-secreting β-cells and 30% glucagon (GCG)-producing α-cells, with the remaining 10% comprised of somatostatin-secreting δ-cells, pancreatic polypeptide (PP) cells, and ghrelin-producing ε-cells^10,15^. While it has been noted that islet distribution varies substantially across the pancreas^12,16^ and small islets tend to have a lower proportion of non–β-cells^17^, we recently reported that islet location within the organ (i.e., head, body, tail) does not appear to influence their endocrine function^9^.

Apart from normal physiology, an improved quantitative and qualitative characterization of human islets is important for understanding the pathogenesis and natural history of many pancreas-based endocrine disorders, including type 1 diabetes (T1D), which results from chronic autoimmunity and the selective loss of INS-producing pancreatic β-cells^18^. In T1D, β-cell loss is preceded by the development of autoantibodies that target β-cell antigens (i.e., glutamic acid decarboxylase (GADA), insulin (IAA), zinc transporter 8 (ZnT8A), and insulinoma associated protein-2 (IA-2A), which typically emerge months to years before clinical symptoms offering valuable prognostic insights^19^. To improve the definition of T1D risk, a disease staging system has been developed that encompasses the number of autoantibodies together with functional assessments of dysglycemia^20^; two or more autoantibodies (stage 1), multiple autoantibodies together with dysglycemia (stage 2), and traditional diagnosis of disease (stage 3 T1D). Though subject to debate, a new category has recently been proposed for persons with a single islet autoantibody and no dysglycemia (stage 0)^21^. While the 10-year risk for T1D development is significantly lower in stage 0 versus stage 1 or 2 T1D, a growing body of evidence suggests that the natural history of T1D is characterized by a series of pancreatic islet abnormalities, including both β- and α-cells, in single GADA+ persons^18^. Understanding the biological processes driving T1D progression, particularly during the earliest stages of disease, will be essential to guide optimal development of therapeutic strategies that preserve endogenous β-cells.

For decades, it was widely suggested that symptomatic disease presents upon 85-95% β-cell loss^22^, yet more recent efforts utilizing tissues obtained from organ donors with recent-onset T1D suggest variability in the residual β-cell mass according to age of disease onset^3,23,24^. Such efforts examining human tissues, while informative, do nonetheless have analytical limitations in terms of their dependence on dimensionality, with most studies utilizing 2D imaging techniques (e.g., hematoxylin and eosin (H&E), immunohistochemistry (IHC), immunofluorescence (IF), imaging mass cytometry (IMC)) to analyze spatial features, rendering complete islet characterization a challenge. Recent advances in 3D imaging techniques have dramatically improved our ability to define essential islet features, including cellular composition/phenotype, size/volume, and spatial distribution within the pancreas^13,14,25,26^. A detailed and comprehensive 3D understanding of normal pancreatic anatomy, along with novel identification of the structural changes that occur in the natural history of T1D, would offer valuable insights into the disease’s development^27–30^.

In this present study, we utilized light sheet fluorescent microscopy (LSFM) to map the 3D distribution of islets in normal human pancreas, obtained from organ donors across adolescence and early adulthood and compared against age-matched individuals with short-duration T1D as well as individuals without diabetes at increased risk for T1D (GADA+, stage 0). These data unveil novel features of β-cell loss during the disorder’s natural history.

## RESULTS AND DISCUSSION

### Decreased islet density and INS+GCG-islet fraction in T1D

We applied LSFM to a collection of transplant grade (i.e., not autopsy) pancreatic tissues (tail region) obtained from organ donors and stained for the endocrine hormones INS and GCG. This age and sex balanced cohort included eight donors with no diabetes (ND), five persons with a single islet autoantibody (GADA+, considered stage 0), and six individuals with short-duration T1D (0-3 years since diagnosis) with residual insulin, representative of T1D’s natural history (Table S1). In the present study, rather than repeatedly utilizing the terms “islets” and “cell clusters”, we have elected to use the term endocrine objects (EO)^31^ to collectively refer to all cell clusters and islet-sized structures, regardless of their hormone expression profile.

As shown, first in representative images of both immunofluorescence (IF) staining (Fig. 1A-I, Fig. S1A-C) and corresponding digital surface renderings for INS and GCG (Fig. 1J-AA, Fig. S1D-F), hormone expression varied dramatically as a function of study group. For this analysis, we assessed volumes of digital surfaces for all INS-containing EO (ICEO, including islets as well as cell clusters; Fig. 1J, M, P), all GCG-containing EO (Fig. 1K, N, Q), as well as INS+GCG-, INS-GCG+, and INS+GCG+ EO displayed all together (Fig. 1L, O, R; Fig. S1D-F) and separately (Fig. 1S-AA).

**Figure 1.**
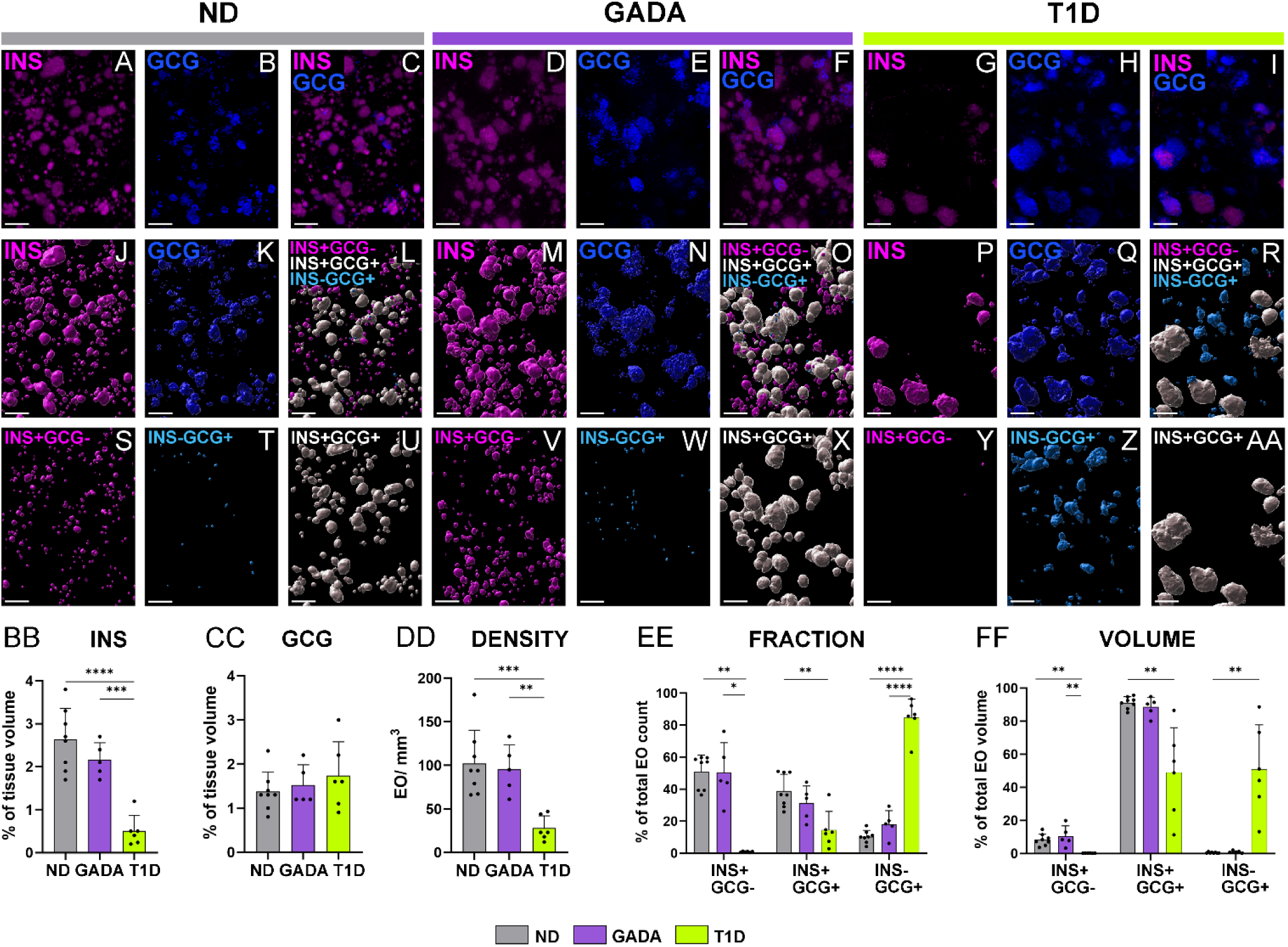
3D quantification of INS+ and GCG+ volume, EO density and cellular composition in human ND, GADA, and T1D pancreas. (A-I) LSFM images show immunofluorescence [IF] staining for (A, D, G) INS in magenta, (B, E, H) GCG in blue, and (C, F, I) overlay in pancreas tail from representative ND [nPOD 6612], GADA+ [nPOD 6582], and T1D [nPOD 6579] donors. (J-AA) Digital surface renderings corresponding to (J, M, P) all INS containing endocrine objects [ICEO], (K, N, Q) all GCG containing EO, and (L, O, R) color-coded EO types shown as an overlay and individually for (S, V, Y) magenta INS+GCG-, (T, W, Z) cyan INS-GCG+, and (U, X, AA) white INS+GCG+. (BB) INS+ and (CC) GCG+ volume normalized as a percentage of the tissue volume. (DD) Total number of EO normalized to the tissue volume. (EE-FF) Islet type expressed as (EE) percentage of total EO count and (FF) percentage of total EO volume. Scale bars are 300 µm. *p<0.05, **p<0.01, ***p < 0.001, ****p < 0.0001; one-way ANOVA or Kruskal-Wallis test, as per methods.

As expected^3^, INS expression was reduced in those with short-duration T1D compared to the ND and GADA+ study groups (Fig. 1BB), with β-cells comprising 0.5±0.3%, 2.6±0.7%, and 2.1±0.3% of the pancreatic volume, respectively. Of note, GADA+ donors were similar to ND in terms of their INS+ tissue volume (Fig. 1BB). α-cell volume was comparable across all three study groups (Fig. 1CC), in line with previous 2D analyses of pancreas sections^3^. Interestingly, however, total EO density (EO/mm³) was lower in the T1D pancreas (27.8±14.3) compared to ND (102.2±38) and GADA+ (95.4±28.1) cases (Fig. 1DD), with approximately 15% of T1D EO containing INS while 85% were INS-deficient (Fig. S1G). Our findings in ND are consistent with a recent study suggesting that INS+ islets constitute approximately 2.8% of the pancreatic volume using LSFM on pancreas tissue from five adult ND donors, along with near-infrared optical projection tomography of an entire human ND pancreas^13^. This said, our data represented investigations of a much wider age range of subjects where age is known to influence islet composition^32,33^. Based on our results (Fig. 1BB-CC), we provide an estimated 2:1 ratio for total INS:GCG content as a percentage of ND pancreatic volume, which further aligns with prior calculations using volumetric estimations from an average islet diameter^12^. Our observations, therefore, dramatically expand on knowledge derived from 2D histological studies of β-cell area and mass before and following T1D onset^4,34,35^.

We noted that in ND and GADA+ pancreata, INS+GCG-EO account for approximately 50% of the total EO count, but only 8–10% of the total EO volume (Fig. 1EE-FF). While perhaps not fitting with historical characterizations of islet endocrine cell content (i.e., 60-70% β cells and 20-30% α-cells (reviewed in ^36^), this observation again corroborates and expands upon a recent analysis from Lehrstrand et al.,^13^ suggesting that approximately 50% of the INS+ EO are virtually devoid of GCG+ α-cells, contributing to nearly 16% of the total islet volume. Indeed, we note INS+GCG+ EO were resident to 38% of the total EO count in ND and 31% in GADA+ (Fig. 1EE), yet represented 91% and 88% of the respective EO volume (Fig. 1FF).

In pancreata obtained from extremely rare, short-duration T1D donors having residual INS content, we noted a dramatic loss of INS+GCG-EO, with additional reductions observed in both INS+GCG+ EO count and volume (Fig. 1EE-FF). In stark contrast with ND and GADA+ persons, INS-GCG+ EO comprised 84% of the T1D EO count (Fig. 1EE) and 50% of the EO volume (Fig. 1FF). We observed that residual INS staining in T1D donors was localized within islets with both hormones present (i.e., INS+GCG+; Fig. 1AA). Recent submissions^2,31^ assessing 2D immunohistochemistry (IHC) stained pancreas sections (Fig. S1H-J), would appear to corroborate these 3D findings. In sum, our 3D analyses suggest that β-cell preservation varies according to EO cell composition.

### INS+ cell clusters and small islets are reduced in the T1D pancreas

Human islet architecture contains diverse patterns of endocrine cell arrangement^37,38^. Figure 2 demonstrates representative distributions of ICEO across entire samples with (Fig. 2A-C) or without shading (Fig. 2D-F) to portray 3D pancreas anatomy within the ND, GADA+, and T1D study groups. ICEO (Fig. 2G-I) and total EO (Fig. 2J-L) were further visualized using a rainbow scale corresponding to EO size. To characterize the differences in hormone content according to EO size, total EO, ICEO, and INS-deficient EO (IDEO) were binned by volume: cell clusters (3,000-10^4^ µm^3^), small islets (10^4^-10^5^ µm^3^), medium islets (10^5^-10^6^ µm^3^), or large islets (≥10^6^ µm^3^). For comparisons with 2D islet imaging data in the literature^11^, these size bins were defined to correspond to the following spherical diameters: 17.9-26.7 µm, 26.8-57.6 µm, 57.7-124.1 µm, and ≥124.1 µm.

**Figure 2.**
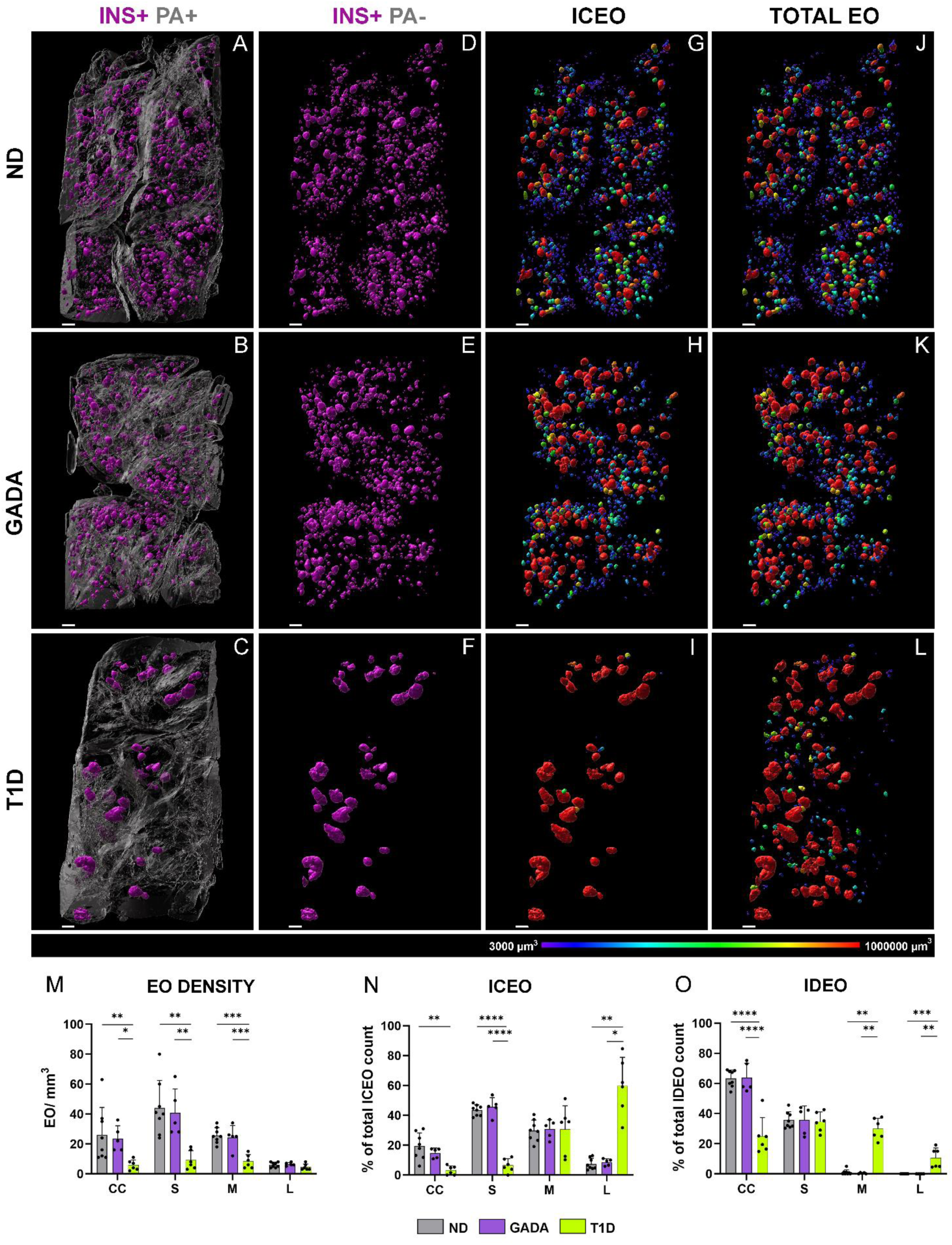
3D islet distribution across human ND, GADA, and T1D pancreas samples. (A-F) Digital surface renderings corresponding to ICEO (A-C) with and (D-F) without pancreas anatomy (PA) shading from representative ND [nPOD 6612], GADA+ [nPOD 6582], and T1D [nPOD 6579] donors. (G-L) Volume color-coded digital surface renderings corresponding to (G-I) ICEO and (J-L) total EO. The heat map line represents the distribution of volumes from 3,000 to 1,000,000 µm^3^. (M) Total number of EO in each size category normalized to the tissue volume. (N-O) Percentage of (N) ICEO and (O) IDEO in each size category. EO size bins: CC – cell clusters, S – small, M – medium, L – large. Scale bars are 300 µm. *p<0.05, **p<0.01, ***p < 0.001, ****p < 0.0001; one-way ANOVA or Kruskal-Wallis test, as per methods.

Overall, EO size densities were similar and normally distributed within the ND and GADA+ groups (Fig. 2M). In comparison, total cell cluster, small- and medium-islet densities were significantly reduced, while large islets were preserved within T1D donors (Fig. 2M). Specifically, the proportions of ICEO were significantly lower within the cell cluster and small-islet size bins, while large islets accounted for the vast majority of ICEO in the T1D group (Fig. 2N). Additionally, IDEO were more frequently observed in T1D (Fig. S1G), but in ND and GADA+ groups, were restricted to cell clusters and small islets (Fig. 2O). Taken together, our findings support the 2D analyses by Seiron et al., demonstrating preservation of islet size despite marked reductions in the total number of islets within the T1D pancreas^39^. However, we suggest a potential alternative interpretation: rather than failure to establish a sufficient number of islets^39^, it remains possible that these data reflect loss of small islets and INS+ cell clusters during T1D development.

### Residual INS in T1D is predominantly preserved in large INS+GCG+ islets

To further explore the endocrine cell composition of islets as a function of T1D natural history, we next analyzed three EO types, defined by the expression of INS and/or GCG, as a function of EO size across the three study groups (Fig. 3). In ND and GADA+ pancreata, INS+GCG-EO, accounting for about half of the total EO count (Fig. 1EE), were preferentially in small size bins, distributed as follows: 16.2±7.4% and 12.7±2.6% in cell clusters, 28.2±5.7% and 29.0±9.8% in small islets, and 6.3±4.2% and 8.7±9.8% in medium islets (Fig. 3A-B). In contrast, the T1D pancreas exhibited a severe reduction in the INS+GCG-EO fraction across these size bins (Fig. 3A-B, Fig. 4A-B). Notably, large INS+GCG- islets were nearly absent across any of the three donor groups (Fig. 3A-B; Fig. 4A-B). In ND and GADA+ donors, INS+GCG+ EO were respectively comprised of 1.1±0.6% and 0.8±0.7% cell clusters, 10.8±5.4% and 6.3±3.8% small islets, 20.4±5.6% and 17.0±8.9% medium islets, and 6.7±3.3% and 7.2±2.3% large islets (Fig. 3A, C). In the ND group, INS+GCG+ EO peaked in the 2x10^5^–4x10^5^ µm^3^ size range, but in the T1D group, the highest density and proportion of total EO count shifted to the 8x10^5^–16x10^5^ µm^3^ range (Fig. 4C-D). Hence, in ND and GADA+ pancreas, INS+GCG+ EO were most commonly identified in the medium-islet size bin, whereas the size of residual INS-containing EO in T1D was primarily shifted to larger INS+GCG+ islets (Fig. 3A, C). INS-GCG+ EO in the ND and GADA+ groups were predominantly represented in the cell cluster and small islet size bins (Fig. 3A, D), with the majority falling below 25x10^3^ µm^3^ (Fig. 4E-F). In the T1D pancreas, INS-GCG+ EO were significantly more prevalent than in ND within all size bins (Fig. 3A, D) and across the continuum of EO size (Fig. 4E-F), likely reflecting the presence of pseudoatrophic islets with irregular or ragged morphology, previously only visualized using 2D microscopy^40–42^. Our ability to rebuild islet boundaries within 3D LSFM images allows for improved visualization of islet morphology beyond these 2D efforts.

**Figure 3.**
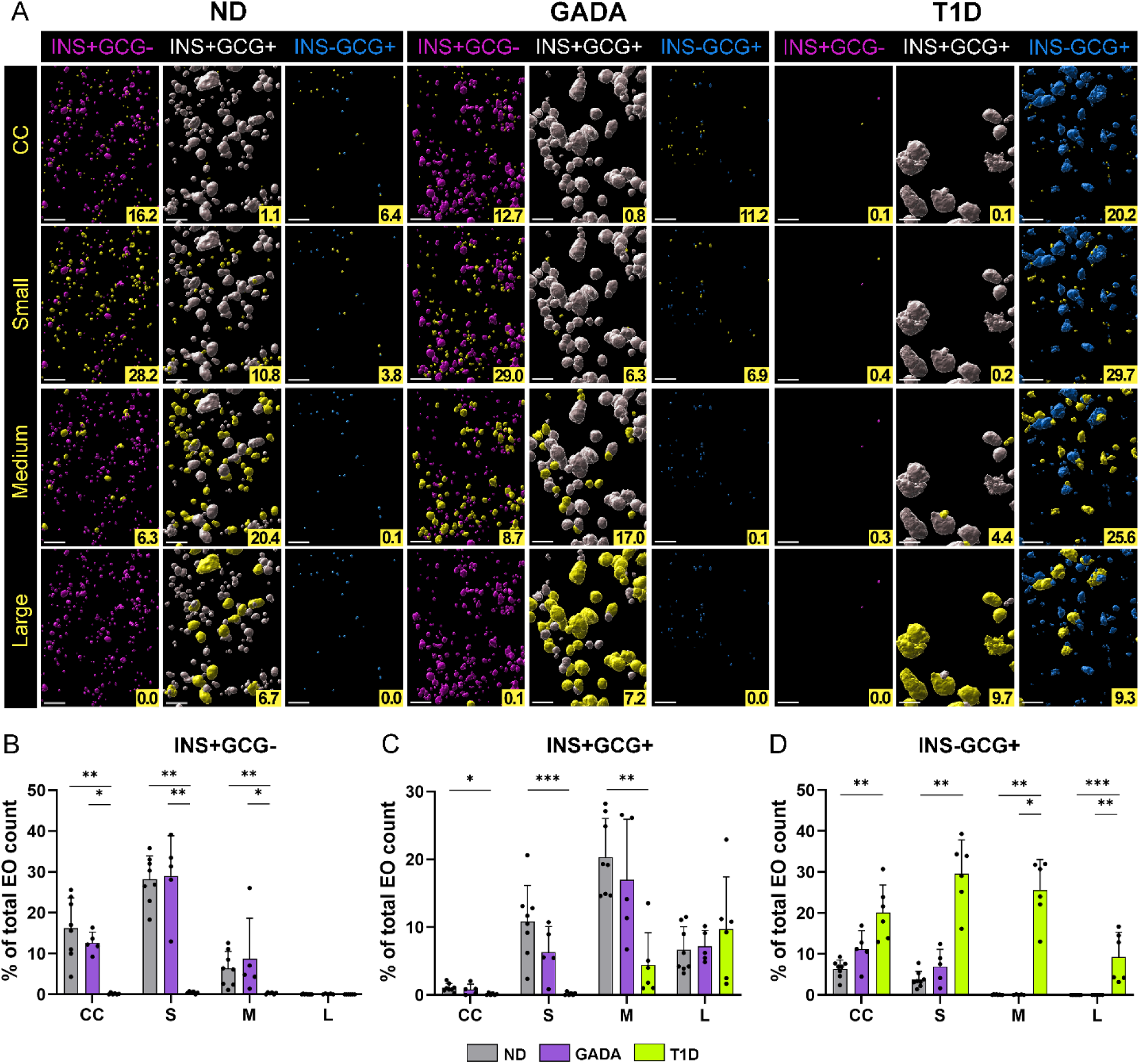
Spatial arrangement and islet size category contribution in human ND, GADA, and T1D pancreas. (A) Digital surface renderings corresponding to color-coded EO types for INS+GCG-(magenta), INS+GCG+ (white), and INS-GCG+ (cyan) with yellow representing each size category in representative ND [nPOD 6612], GADA+ [nPOD 6582], and T1D [nPOD 6579] donors. The numbers in the yellow boxes indicate the percentage of EO in each size category. Scale bars are 300 µm. (B) INS+GCG-, (C) INS+GCG+, and (D) INS-GCG+ EO normalized as a percentage of total EO count. EO size bins: CC – cell clusters, S – small, M – medium, L – large. *p<0.05, **p<0.01, ***p < 0.001, ****p < 0.0001; Kruskal-Wallis test, as per methods.

**Figure 4.**
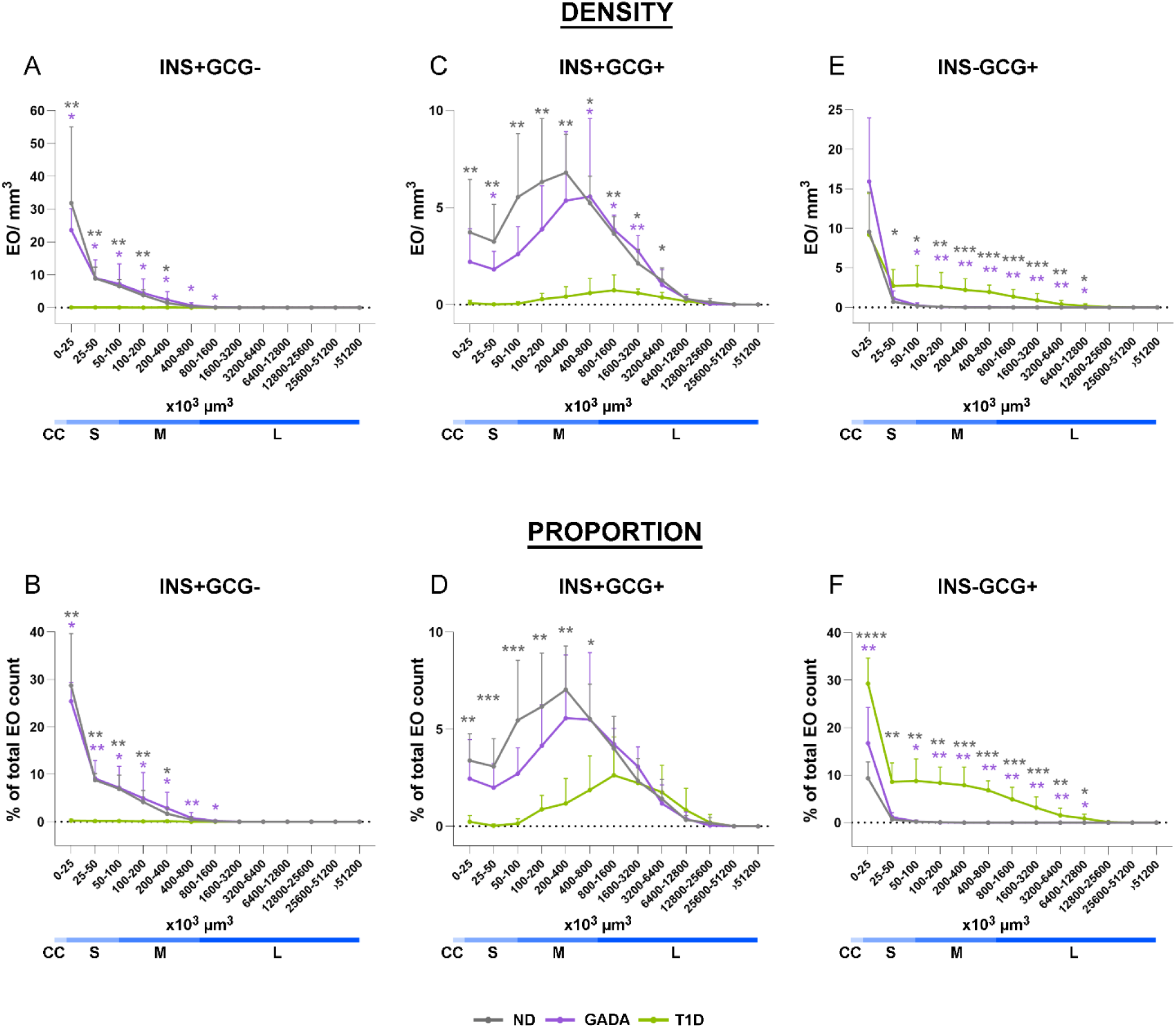
Statistical analysis across the continuous islet size distribution in human ND, GADA, and T1D pancreas. (A, B) INS+GCG-, (C, D) INS+GCG+, and (E, F) INS-GCG+ EO expressed as (A, C, E) normalized to tissue volume and (B, D, F) percentage of total EO count. Gray asterisks show difference between ND and T1D groups, purple asterisks show difference between GADA and T1D groups, ND vs GADA p=ns. *p<0.05, **p<0.01, ***p < 0.001; one-way ANOVA or Kruskal-Wallis test, as per methods.

The significant loss of INS+ cell clusters and small islets in T1D suggests that size represents an important feature in terms of susceptibility to autoimmune destruction. The outstanding question remains as to why this occurs. As medium and large islets contain a more diverse mix of endocrine cells, including higher number of α-cells^13^, this may be an important contributor to their survival. In contrast, small INS+ EO, which are predominantly composed of β-cells, have higher insulin content and are capable of greater insulin secretion as compared to large islets^17,43^. As insulin represents a key autoantigen in T1D^44^, these features may also contribute to greater susceptibility to autoimmune attack and destruction.

The large number of small EO observed in the INS+GCG-fraction within ND pancreas, coupled with their dramatic loss in short-duration T1D, raises critical questions regarding their potential specialization, distinct functionality, and/or susceptibility to destruction as compared with larger INS+GCG+ islets. Taken together with a recent report involving 2D imaging^31^, we expect that the role of INS+ small islets and cell clusters is likely underappreciated in T1D pathogenesis, particularly considering that in younger individuals, who have smaller islets overall^45,46^, T1D progresses more rapidly^47^. In sum, the loss of small INS+GCG- islets and cell clusters represents a characteristic feature of the T1D pancreas, while larger INS+GCG+ islets seem to preferentially persist, suggesting cell composition and size-dependent resilience.

### Limitations of the study

Studies of human pancreas samples suffer from numerous pragmatic challenges. First among these is the extensive effort required to collect high-quality pancreata in large numbers relative to, for example, assessing peripheral blood from living persons, as the human pancreas is not subject to biopsy due to ethical considerations^48^. Yet, through extensive attempts by many over the last two decades, we and others have been successful in obtaining cases to support the numbers in this study. Second, unlike animal models or again, studies of living humans, longitudinal analyses are not feasible from organ donor tissues; hence, interpretations seeking to infer disease pathogenesis and natural history must be made through collation of cross-sectional data. This said, as all tissues were processed consistently in accordance with SOP-driven protocols; hence, we do not see where a bias may have occurred with respect to a given study group. In addition, based on technical practices of U.S. Organ Procurement Organizations, our efforts were not afforded the ability to screen for IAA^49^. Hence, there is no pragmatic means in place to identify this otherwise highly interesting population at increased risk for T1D. Future 3D studies using LSFM are underway to observe changes occurring closer to T1D onset in donors with multiple islet autoantibodies and with the addition of immune markers to examine insulitis during the natural history of the disease.

## RESOURCE AVAILABILITY

### Lead contact

Requests for further information and resources should be directed to the lead contact, Dr. Mark Atkinson.

### Materials availability

Additional tissue samples and whole slide images from donors evaluated as a part of this study can be requested from nPOD for use in projects approved by the nPOD Tissue Prioritization Committee, as outlined on the nPOD website (www.npod.org). This study did not generate new unique reagents.

### Data and code availability

All data and any additional information required to reanalyze the data reported in this paper is available from the lead contact upon request. No original code was utilized for the analyses reported herein.

## ACKNOWLEDGMENTS

We thank the organ donors and their families. This study was supported by the following grants from the National Institutes of Health (NIH): R01DK131059, R01DK123292, and P01AI042288 to MAA, U24DK104162 to JSK, and U01DK135001 to CHW. This research was performed with the support of the Network for Pancreatic Organ donors with Diabetes (nPOD; RRID:SCR_014641), a collaborative type 1 diabetes research project supported by Breakthrough T1D and The Leona M. & Harry B. Helmsley Charitable Trust (Grant# 3-SRA-2023-1417-S-B). The content and views expressed are the responsibility of the authors and do not necessarily reflect those of nPOD. Organ Procurement Organizations (OPO) partnering with nPOD to provide research resources are listed at https://npod.org/for-partners/npod-partners/.

## AUTHOR CONTRIBUTIONS

Conceptualization: M.A.A.; Supervision: A.L.P., M.A.A.; Resources: I.K., M.A.A., Funding acquisition: C.H.W., M.A.A.; Data Curation: A.R., S.C.; Formal Analysis: A.R.; Statistical calculations: A.R. and J.S.K.; Investigation: A.R., A.L.P., M.B., C.H.W., M.C.T., M.A.A.; Methodology: A.R., S.C.; Project Administration: A.L.P., I.K., M.D.W.; Writing: Original draft A.R., A.L.P., M.A.A.; Writing: Review & Editing: A.R., A.L.P., M.D.W., M.B., C.H.W., J.S.K., I.K., M.C.T., M.A.A.

## DECLARATION OF INTERESTS

The authors declare no competing interests.

## SUPPLEMENTAL INFORMATION

**Table S1. Donor information.**

**Figures S1.**
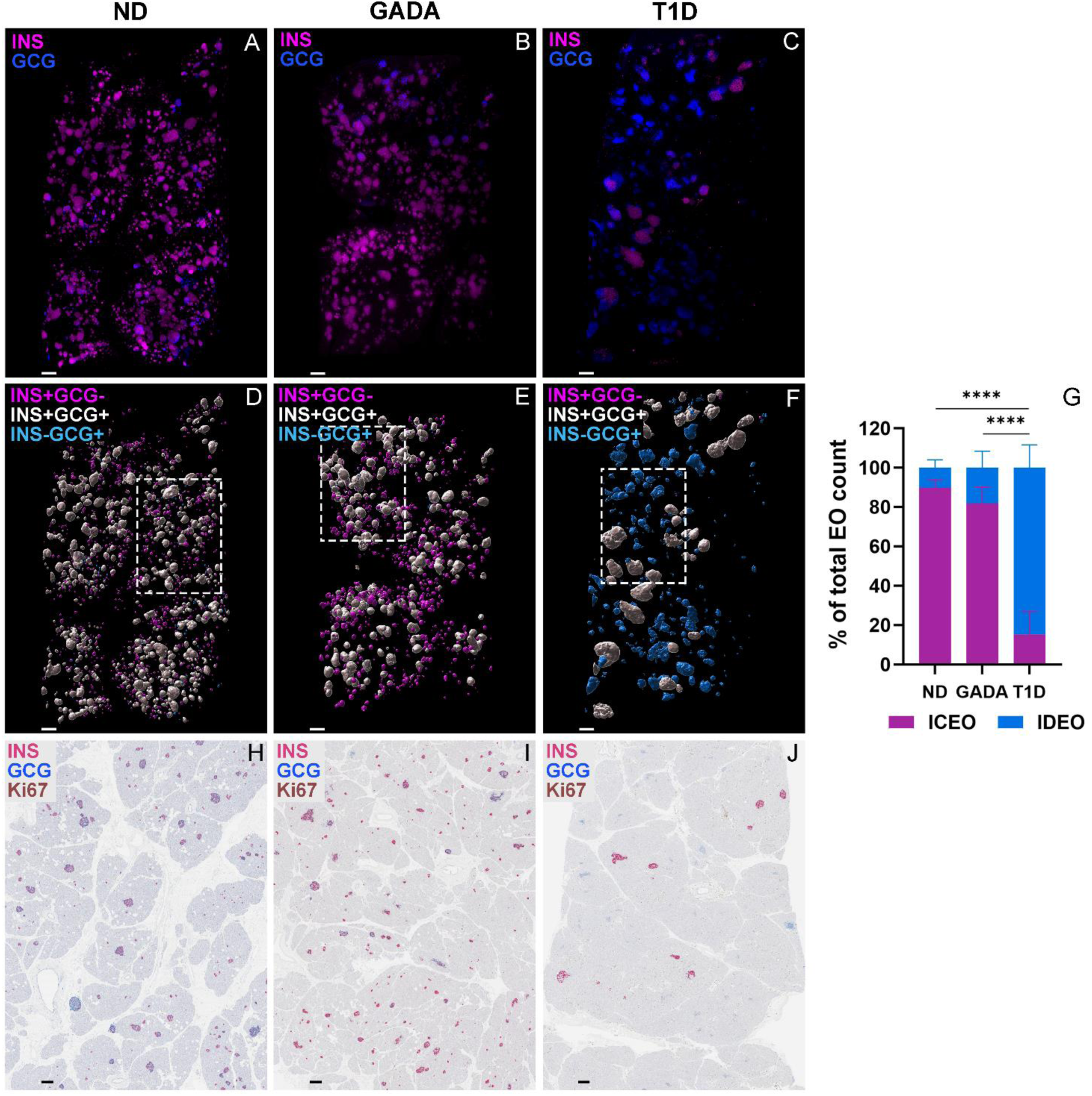
Spatial arrangement of islet type categories across samples in ND, GADA and T1D pancreas. (A-C) LSFM image overlays of INS+ (magenta) and GCG+ (blue) in pancreas tail from representative ND [nPOD 6612], GADA+ [nPOD 6582], and T1D [nPOD 6579] donors. (D-F) Digital surface renderings for color-coded endocrine object (EO) types shown as overlays of INS+GCG-(magenta), INS+GCG+ (white), and INS-GCG+ (cyan) with dashed lines around the region of interest shown in Fig. 1L, O, R. (G) INS-containing endocrine objects (ICEO) and INS-deficient endocrine objects (IDEO) normalized as a percentage of total EO count, ****p < 0.0001, one-way ANOVA. (H-J) INS (pink), GCG (blue), and Ki67 (brown) immunohistochemistry (IHC) images from the nPOD Aperio digital pathology database for the same three donors. All scale bars are 300 mm.

## STAR★METHODS

### KEY RESOURCES TABLE

| REAGENT or RESOURCE | SOURCE | IDENTIFIER |
| --- | --- | --- |
| Antibodies |  |  |
| Monoclonal Rabbit Anti-Insulin | Abcam | Cat# ab181547; RRID: AB_2716761 |
| Monoclonal Rabbit Anti-Glucagon conjugated to Alexa Fluor® 488 | Abcam | Cat# ab307340 |
| Biological samples |  |  |
| Human Pancreas Tissue | nPOD;https://npod.org/ | RRID: SCR_014641 |
| nPOD-6550 |  | RRID: SAMN25652261 |
| nPOD-6566 |  | RRID: SAMN33284286 |
| nPOD-6569 |  | RRID: SAMN38117299 |
| nPOD-6573 |  | RRID: SAMN33284291 |
| nPOD-6575 |  | RRID: SAMN33284293 |
| nPOD-6578 |  | RRID: SAMN33284295 |
| nPOD-6579 |  | RRID: SAMN33284296 |
| nPOD-6582 |  | RRID: SAMN38117302 |
| nPOD-6588 |  | RRID: SAMN40555580 |
| nPOD-6593 |  | RRID: SAMN38117312 |
| nPOD-6596 |  | RRID: SAMN40555583 |
| nPOD-6598 |  | RRID: SAMN40555585 |
| nPOD-6601 |  | RRID: SAMN40555588 |
| nPOD-6604 |  | RRID: SAMN40555591 |
| nPOD-6605 |  | RRID: SAMN40555592 |
| nPOD-6607 |  | RRID: SAMN40555594 |
| nPOD-6611 |  | RRID: SAMN44486497 |
| nPOD-6612 |  | RRID: SAMN44486498 |
| nPOD-6615 |  | RRID: SAMN44486501 |
| Chemicals, peptides, and recombinant proteins |  |  |
| Ethyl cinnamate | MilliporeSigma | Cat# W243000-1KG-K |
| Triton X-100 | MilliporeSigma | Cat# 648462 |
| 4% Paraformaldehyde | Thermo Fisher Scientific | Cat# J19943K2 |
| Dimethyl Sulfoxide | Thermo Fisher Scientific | Cat# D128-500 |
| Urea | Sigma-Aldrich | Cat# U5128-1KG |
| Glycerol | Thermo Scientific | Cat# 17904 |
| PBS with Azide | Santa Cruz Biotechnology | Cat# SC-296028 |
| 10X PBS buffer | Invitrogen | Cat# AM9625 |
| Animal-Free Blocker (5x) concentrate | Vector Laboratories | Cat# SP5030 |
| Tween 20 | MilliporeSigma | Cat# P6585 |
| Methanol | Fisher Chemical | Cat# A412-4 |
| Dibenzyl ether | MilliporeSigma | Cat# 33630-1L |
| Dichloromethane | MilliporeSigma | Cat# 270997-2L |
| Critical commercial assays |  |  |
| Alexa Fluor™ Antibody Labeling Kit 784/814 nm | Molecular Probes | Cat# A20189 |
| Software and algorithms |  |  |
| Prism 10 | GraphPad | RRID: SCR_002798 |
| ImSpector Pro version 7.6.3.0 | LaVision BioTec | N/A |
| Imaris 10.2 | Oxford Instruments | RRID: SCR_007370 |
| Other |  |  |
| Centrifuge 5424R | Eppendorf | SN 5404IQ239594 |
| MaxQ™ 4450 Orbital Shaker | Thermo Fisher Scientific | Model SHKE 4450 |
| VWR Rocking Platform Shakers, 230 V | VWR international | Model 10860-660 |
| Ultra Microscope Blaze™ | Miltenyi Biotec | Cat# 150-001-760 |

### EXPERIMENTAL MODEL AND STUDY PARTICIPANT DETAILS

Pancreas tissues from human donors with or without T1D were procured with informed consent provided by each donor’s legal representative as approved by the United Network for Organ Sharing (UNOS) and according to federal guidelines. Donor pancreata were recovered, placed in transport media on ice, and shipped via organ courier to the nPOD program at the University of Florida (RRID: SCR_014641, https://www.jdrfnpod.org). The tissue was processed by a licensed pathology assistant according to the nPOD Organ Processing and Pathology Core (OPPC) standard operating procedures approved by the University of Florida Institutional Review Board (IRB201600029). For each donor, a medical chart review was performed and C-peptide measured to confirm T1D diagnosis according to guidelines established by the American Diabetes Association (ADA). Demographic data, hospitalization duration, and organ transport time were obtained from hospital records. Donor information is listed in Table S1.

## METHOD DETAILS

### Pancreas Sample Preparation

Samples were freshly harvested from the pancreas tail region (0.7-1cm^3^ in size) and fixed in 4% PFA [Thermo Scientific] for 4 days at 4°C while shaking at 80 rotations per minute (rpm) [VWR Rocking Platform Shakers, 230 V]. Fixed samples were washed with 1X PBS [Invitrogen] for 1 day at 37°C while shaking at 80 rpm [Thermo Scientific MaxQ™ 4450 Orbital Shaker], then stored in 1X PBS with azide [Santa Cruz] at 4°C.

### 3D Immunostaining

Using modified iDISCO protocol^50^ fixed human pancreas samples (approximately X=5mm, Y=10mm, Z=3mm) were stepwise dehydrated for 1 hour at 37°C with shaking at 80 rpm in increasing concentrations of methanol (MeOH) [Fisher Chemical] in deionized water (diH_2_O; 20%, 40%, 60%, 80%, 100%). Samples were photobleached in 5% H_2_O_2_ in MeOH for 4 days at 4°C under LED light, then washed in 100% MeOH for 1 hour at room temperature (RT) while shaking at 80 rpm and then, stored at 4°C. For delipidating, samples were incubated overnight at RT in a 2:1 solution of dichloromethane (DCM) [Sigma-Aldrich] and MeOH, then at RT shaking 80 rpm in 100% DCM for 1 hour and washed in 100% MeOH two times for 30 minutes. Next, samples were re-hydrated for 1 hour at 37°C while shaking 80 rpm in decreasing concentrations of MeOH in diH_2_O (80%, 60%, 40%, 20%), then washed in 1X PBS two times. Permeabilization was performed overnight at 37°C 80 rpm in permeabilization buffer (25% urea [Sigma-Aldrich], 15% glycerol [Thermo Scientific], 7% Triton X-100 [Sigma-Aldrich] in 1X PBS). Then, the samples were washed five times for 1 hour at 37°C while shaking 80 rpm in 1X PBS. Blocking was performed overnight at 37°C with shaking 80 rpm in blocking buffer (10% animal-free blocker [Vector Laboratories], 10% DMSO [Fisher Chemical], 0.5% Triton X-100 [Sigma-Aldrich], 0.2% Tween 20 [Sigma-Aldrich] in 1X PBS).

The anti-human INS primary antibody (Cat. No. ab181547; Abcam) was conjugated with Alexa Fluor 790 dye using the Alexa Fluor™ Antibody Labeling Kit (Cat. No. A20189; Molecular Probes™) according to the manufacturer’s protocol. For whole-mount immunostaining, samples were incubated with conjugated primary antibodies specific for GCG (Alexa Fluor® 488 anti-human GCG; Cat. No. ab307340; Abcam; diluted 1:5000) and INS (anti-human INS Alexa Fluor 790; diluted 1:400) diluted in a staining buffer (1X PBS with azide containing 2% animal-free blocker, 10% DMSO, 0.5% Triton X-100). The master mix of antibodies was centrifuged at 15,000g for 15 minutes to prevent the formation of fluorophore precipitates in samples, then applied to samples and incubated at 37°C for 6 days while shaking at 80 rpm. After antibody labeling, all samples were washed for 1 hour at 37°C while shaking 80 rpm, four times in wash buffer (2% animal-free blocker [Vector Laboratories], 0.5% Triton X-100 [Sigma-Aldrich] in 1X PBS) and one time in 1X PBS. Next, samples were stepwise dehydrated for 1 hour at 37°C while shaking at 80 rpm in increasing concentrations of MeOH (20%, 40%, 60%, 80%, 100%, 100%). Samples were stored in 100% MeOH in the dark at RT until clearing.

### 3D Tissue Clearing

Samples were incubated in a 2:1 solution of DCM and MeOH for 1-3 hours at RT with shaking 80 rpm until the sample sank to the bottom of the tube, followed by two 15-minute incubations in 100% DCM. Then, the samples were transferred to dibenzyl ether (DBE) [Sigma-Aldrich] to clear for 3-24 hours. Samples were stored in DBE for 72 hours to avoid a decrease in signal intensity.

### 3D Light Sheet Fluorescence Microscopy (LSFM) Imaging

Image acquisition was performed with an Ultra Microscope Blaze (Miltenyi Biotec, Germany) and ImSpector Pro (version 7.6.3.0, LaVision BioTec, Germany) acquisition software. Laser light sheets were generated at excitation wavelengths of 488, 640, and 785 nm using an objective lens with 4× magnification (MI Plan 4× NA0.35). All samples were scanned at 1× zoom magnification using ImSpector Pro acquisition software. The *z*-step size was 2 μm. Filter sets used were as follows: GCG, excitation (Ex): 488, emission (Em): 525/50; autofluorescence (pancreas anatomy), Ex: 640, Em: 680/30; and INS, Ex: 785, Em: 845/55. The resultant image datasets were saved in *ome.tif format.

## QUANTIFICATION AND STATISTICAL ANALYSIS

### 3D Reconstruction and Surfacing

Image *ome.tif files were converted into *ims files using Imaris File Converter software (version 10.2.0; Bitplane, UK). Imaris software (version 10.2.0; Bitplane, UK) was used to create 3D images and then, surfaces for pancreas anatomy, GCG+, and INS+ signals. Surfaces for both high-intensity and low-intensity objects were generated. The INS surfaces for low-intensity objects were added to those for high-intensity objects with a filter applied to exclude overlapping objects with a minimum overlap of 10 μm^3^, and the two resultant surfaces were merged to account for objects including an intensity gradient of INS signal. To accurately cover islets with both GCG+ and INS+ signals, the machine learning option was applied in Imaris to generate entire islet surfaces. INS-GCG+ surfaces on image borders were excluded from quantification to account for only true INS-GCG+ EO. The manual section-by-section quality control in the Imaris slice scanner was applied to confirm the detection of INS and/or GCG signal presence in islets. Artifacts outside the tissue volume surface (determined from autofluorescence) were also manually excluded.

### Statistics

EO with volume ≥3000µm^3^ were binned by volume (endocrine cell clusters: 3x10^3^-10^4^, small: 10^4^-10^5^, medium: 10^5^-10^6^, and large islets: ≥10^6^ µm^3^). A one-way Analysis of Variance (ANOVA) was used to examine the differences in tissue or EO volume, density, and/or count by donor type. The assumptions of ANOVA were examined using the Shapiro-Wilk test for normality, and a Bartlett’s test for homogeneity of variance (i.e. heteroscedasticity); all data are independent. If the assumptions held, a one-way ANOVA was used together with a post-hoc Bonferroni’s test for multiple comparisons. If the assumptions were not met, data were analyzed by Kruskal-Wallis test followed by a post-hoc Dunn’s test for multiple comparisons. All results are presented as mean ± standard deviation (SD). p-values less than the nominal alpha level of 0.05 indicate statistical significance. All analyses were performed using GraphPad Prism software version 10.

## ADDITIONAL RESOURCES

The nPOD Data Portal contains detailed information about each donor tissue used in this study: https://portal.jdrfnpod.org/.

## Notes

### Competing Interest Statement

The authors have declared no competing interest.

## REFERENCES

1. Atkinson, M.A., Campbell-Thompson, M., Kusmartseva, I., and Kaestner, K.H. (2020). Organisation of the human pancreas in health and in diabetes. Diabetologia 63, 1966–1973. 10.1007/s00125-020-05203-7.

2. van der Heide, V., McArdle, S., Nelson, M.S., Cerosaletti, K., Gnjatic, S., Mikulski, Z., Posgai, A.L., Kusmartseva, I., Atkinson, M., and Homann, D. (2025). Integrated histopathology of the human pancreas throughout stages of type 1 diabetes progression. bioRxiv. 10.1101/2025.03.18.644000.

3. Campbell-Thompson, M., Fu, A., Kaddis, J.S., Wasserfall, C., Schatz, D.A., Pugliese, A., and Atkinson, M.A. (2016). Insulitis and beta-Cell Mass in the Natural History of Type 1 Diabetes. Diabetes 65, 719–731. 10.2337/db15-0779.

4. Rodriguez-Calvo, T., Zapardiel-Gonzalo, J., Amirian, N., Castillo, E., Lajevardi, Y., Krogvold, L., Dahl-Jorgensen, K., and von Herrath, M.G. (2017). Increase in Pancreatic Proinsulin and Preservation of Beta Cell Mass in Autoantibody Positive Donors prior to Type 1 Diabetes Onset. Diabetes. 10.2337/db16-1343.

5. Saisho, Y., Butler, A.E., Manesso, E., Elashoff, D., Rizza, R.A., and Butler, P.C. (2013). beta-cell mass and turnover in humans: effects of obesity and aging. Diabetes Care 36, 111–117. 10.2337/dc12-0421.

6. Espes, D., Manell, E., Ryden, A., Carlbom, L., Weis, J., Jensen-Waern, M., Jansson, L., and Eriksson, O. (2019). Pancreatic perfusion and its response to glucose as measured by simultaneous PET/MRI. Acta diabetologica 56, 1113–1120. 10.1007/s00592-019-01353-2.

7. Fujimoto, H., Fujita, N., Hamamatsu, K., Murakami, T., Nakamoto, Y., Saga, T., Ishimori, T., Shimizu, Y., Watanabe, H., Sano, K., et al. (2021). First-in-Human Evaluation of Positron Emission Tomography/Computed Tomography With [(18)F]FB(ePEG12)12-Exendin-4: A Phase 1 Clinical Study Targeting GLP-1 Receptor Expression Cells in Pancreas. Front Endocrinol (Lausanne) 12, 717101. 10.3389/fendo.2021.717101.

8. Hart, N.J., and Powers, A.C. (2019). Use of human islets to understand islet biology and diabetes: progress, challenges and suggestions. Diabetologia 62, 212–222. 10.1007/s00125-018-4772-2.

9. Drotar, D.M., Mojica-Avila, A.K., Bloss, D.T., Cohrs, C.M., Manson, C.T., Posgai, A.L., Williams, M.D., Brusko, M.A., Phelps, E.A., Wasserfall, C.H., et al. (2024). Impaired islet function and normal exocrine enzyme secretion occur with low inter-regional variation in type 1 diabetes. Cell Rep 43, 114346. 10.1016/j.celrep.2024.114346.

10. Da Silva Xavier, G. (2018). The Cells of the Islets of Langerhans. J Clin Med 7. 10.3390/jcm7030054.

11. Huang, H.H., Harrington, S., and Stehno-Bittel, L. (2018). The Flaws and Future of Islet Volume Measurements. Cell Transplant 27, 1017–1026. 10.1177/0963689718779898.

12. Ionescu-Tirgoviste, C., Gagniuc, P.A., Gubceac, E., Mardare, L., Popescu, I., Dima, S., and Militaru, M. (2015). A 3D map of the islet routes throughout the healthy human pancreas. Scientific reports 5, 14634. 10.1038/srep14634.

13. Lehrstrand, J., Davies, W.I.L., Hahn, M., Korsgren, O., Alanentalo, T., and Ahlgren, U. (2024). Illuminating the complete ss-cell mass of the human pancreas-signifying a new view on the islets of Langerhans. Nature communications 15, 3318. 10.1038/s41467-024-47686-7.

14. Hahn, M., Nord, C., Eriksson, M., Morini, F., Alanentalo, T., Korsgren, O., and Ahlgren, U. (2021). 3D imaging of human organs with micrometer resolution - applied to the endocrine pancreas. Commun Biol 4, 1063. 10.1038/s42003-021-02589-x.

15. Dolensek, J., Rupnik, M.S., and Stozer, A. (2015). Structural similarities and differences between the human and the mouse pancreas. Islets 7, e1024405. 10.1080/19382014.2015.1024405.

16. Ravi, P.K., Singh, S.R., and Mishra, P.R. (2021). Redefining the tail of pancreas based on the islets microarchitecture and inter-islet distance: An immunohistochemical study. Medicine 100, e25642. 10.1097/MD.0000000000025642.

17. Dybala, M.P., and Hara, M. (2019). Heterogeneity of the Human Pancreatic Islet. Diabetes 68, 1230–1239. 10.2337/db19-0072.

18. Atkinson, M.A., and Mirmira, R.G. (2023). The pathogenic "symphony" in type 1 diabetes: A disorder of the immune system, β cells, and exocrine pancreas. Cell Metabolism 35, 1500–1518. 10.1016/j.cmet.2023.06.018.

19. Felton, J.L., Redondo, M.J., Oram, R.A., Speake, C., Long, S.A., Onengut-Gumuscu, S., Rich, S.S., Monaco, G.S.F., Harris-Kawano, A., Perez, D., et al. (2024). Islet autoantibodies as precision diagnostic tools to characterize heterogeneity in type 1 diabetes: a systematic review. Commun Med (Lond) 4, 66. 10.1038/s43856-024-00478-y.

20. Insel, R.A., Dunne, J.L., Atkinson, M.A., Chiang, J.L., Dabelea, D., Gottlieb, P.A., Greenbaum, C.J., Herold, K.C., Krischer, J.P., Lernmark, A., et al. (2015). Staging presymptomatic type 1 diabetes: a scientific statement of JDRF, the Endocrine Society, and the American Diabetes Association. Diabetes Care 38, 1964–1974. 10.2337/dc15-1419.

21. Manescu, M., Manescu, I.B., and Grama, A. (2024). A Review of Stage 0 Biomarkers in Type 1 Diabetes: The Holy Grail of Early Detection and Prevention? J Pers Med 14. 10.3390/jpm14080878.

22. Battaglia, M., and Atkinson, M.A. (2015). The streetlight effect in type 1 diabetes. Diabetes 64, 1081–1090. 10.2337/db14-1208.

23. Damond, N., Engler, S., Zanotelli, V.R.T., Schapiro, D., Wasserfall, C.H., Kusmartseva, I., Nick, H.S., Thorel, F., Herrera, P.L., Atkinson, M.A., and Bodenmiller, B. (2019). A Map of Human Type 1 Diabetes Progression by Imaging Mass Cytometry. Cell Metab 29, 755–768.e755. 10.1016/j.cmet.2018.11.014.

24. Rodriguez-Calvo, T., Richardson, S.J., and Pugliese, A. (2018). Pancreas Pathology During the Natural History of Type 1 Diabetes. Curr Diab Rep 18, 124. 10.1007/s11892-018-1084-3.

25. Hahn, M., Nord, C., Franklin, O., Alanentalo, T., Mettavainio, M.I., Morini, F., Eriksson, M., Korsgren, O., Sund, M., and Ahlgren, U. (2020). Mesoscopic 3D imaging of pancreatic cancer and Langerhans islets based on tissue autofluorescence. Scientific reports 10, 18246. 10.1038/s41598-020-74616-6.

26. Villalba, A., Gitton, Y., Inoue, M., Aiello, V., Blain, R., Toupin, M., Mazaud-Guittot, S., Rachdi, L., Semb, H., Chedotal, A., and Scharfmann, R. (2024). A 3D atlas of the human developing pancreas to explore progenitor proliferation and differentiation. Diabetologia 67, 1066–1078. 10.1007/s00125-024-06143-2.

27. Campbell-Thompson, M., and Tang, S.C. (2021). Pancreas Optical Clearing and 3-D Microscopy in Health and Diabetes. Frontiers in Endocrinology (Lausanne) 12, 644826. 10.3389/fendo.2021.644826.

28. Alvarsson, A., Jimenez-Gonzalez, M., Li, R., Rosselot, C., Tzavaras, N., Wu, Z., Stewart, A.F., Garcia-Ocana, A., and Stanley, S.A. (2020). A 3D atlas of the dynamic and regional variation of pancreatic innervation in diabetes. Sci Adv 6. 10.1126/sciadv.aaz9124.

29. Campbell-Thompson, M., Butterworth, E.A., Boatwright, J.L., Nair, M.A., Nasif, L.H., Nasif, K., Revell, A.Y., Riva, A., Mathews, C.E., Gerling, I.C., et al. (2021). Islet sympathetic innervation and islet neuropathology in patients with type 1 diabetes. Scientific reports 11, 6562. 10.1038/s41598-021-85659-8.

30. Hsueh, B., Burns, V.M., Pauerstein, P., Holzem, K., Ye, L., Engberg, K., Wang, A.C., Gu, X., Chakravarthy, H., Arda, H.E., et al. (2017). Pathways to clinical CLARITY: volumetric analysis of irregular, soft, and heterogeneous tissues in development and disease. Scientific reports 7, 5899. 10.1038/s41598-017-05614-4.

31. Murrall, K., Luckett, T., Lekka, C., Flaxman, C.S., Wyatt, R., Akhbari, P., Kusmartseva, I., Hunter, S.L., Leete, P., Burn, I., et al. (2025). Small things matter: Lack of extra-islet beta cells in Type 1 diabetes. bioRxiv. 10.1101/2025.04.11.648319.

32. Arda, H.E., Li, L., Tsai, J., Torre, E.A., Rosli, Y., Peiris, H., Spitale, R.C., Dai, C., Gu, X., Qu, K., et al. (2016). Age-Dependent Pancreatic Gene Regulation Reveals Mechanisms Governing Human beta Cell Function. Cell Metab 23, 909–920. 10.1016/j.cmet.2016.04.002.

33. Shrestha, S., Erikson, G., Lyon, J., Spigelman, A.F., Bautista, A., Manning Fox, J.E., Dos Santos, C., Shokhirev, M., Cartailler, J.P., Hetzer, M.W., et al. (2022). Aging compromises human islet beta cell function and identity by decreasing transcription factor activity and inducing ER stress. Sci Adv 8, eabo3932. 10.1126/sciadv.abo3932.

34. Diedisheim, M., Mallone, R., Boitard, C., and Larger, E. (2016). Beta-cell mass in non-diabetic autoantibody-positive subjects: an analysis based on the nPOD database. J Clin Endocrinol Metab, jc20153756. 10.1210/jc.2015-3756.

35. In’t Veld, P., Lievens, D., De Grijse, J., Ling, Z., Van der Auwera, B., Pipeleers-Marichal, M., Gorus, F., and Pipeleers, D. (2007). Screening for insulitis in adult autoantibody-positive organ donors. Diabetes 56, 2400–2404. db07-0416 [pii] 10.2337/db07-0416.

36. Walker, J.T., Saunders, D.C., Brissova, M., and Powers, A.C. (2021). The Human Islet: Mini-Organ With Mega-Impact. Endocr Rev 42, 605–657. 10.1210/endrev/bnab010.

37. Brissova, M., Fowler, M.J., Nicholson, W.E., Chu, A., Hirshberg, B., Harlan, D.M., and Powers, A.C. (2005). Assessment of human pancreatic islet architecture and composition by laser scanning confocal microscopy. J Histochem Cytochem 53, 1087–1097. 10.1369/jhc.5C6684.2005.

38. Steiner, D.J., Kim, A., Miller, K., and Hara, M. (2010). Pancreatic islet plasticity: interspecies comparison of islet architecture and composition. Islets 2, 135–145. 10.4161/isl.2.3.11815.

39. Seiron, P., Wiberg, A., Kuric, E., Krogvold, L., Jahnsen, F.L., Dahl-Jorgensen, K., Skog, O., and Korsgren, O. (2019). Characterisation of the endocrine pancreas in type 1 diabetes: islet size is maintained but islet number is markedly reduced. J Pathol Clin Res 5, 248–255. 10.1002/cjp2.140.

40. Campbell-Thompson, M.L., Atkinson, M.A., Butler, A.E., Chapman, N.M., Frisk, G., Gianani, R., Giepmans, B.N., von Herrath, M.G., Hyoty, H., Kay, T.W., et al. (2013). The diagnosis of insulitis in human type 1 diabetes. Diabetologia 56, 2541–2543. 10.1007/s00125-013-3043-5.

41. Gepts, W. (1965). Pathologic anatomy of the pancreas in juvenile diabetes mellitus. Diabetes 14, 619–633.

42. Wang, Y.J., Traum, D., Schug, J., Gao, L., Liu, C., Atkinson, M.A., Powers, A.C., Feldman, M.D., Naji, A., Chang, K.M., and Kaestner, K.H. (2019). Multiplexed In Situ Imaging Mass Cytometry Analysis of the Human Endocrine Pancreas and Immune System in Type 1 Diabetes. Cell Metab 29, 769–783.e764. 10.1016/j.cmet.2019.01.003.

43. Farhat, B., Almelkar, A., Ramachandran, K., Williams, S.J., Huang, H.H., Zamierowksi, D., Novikova, L., and Stehno-Bittel, L. (2013). Small human islets comprised of more beta-cells with higher insulin content than large islets. Islets 5, 87–94. 10.4161/isl.24780.

44. Kent, S.C., Mannering, S.I., Michels, A.W., and Babon, J.A.B. (2017). Deciphering the Pathogenesis of Human Type 1 Diabetes (T1D) by Interrogating T Cells from the "Scene of the Crime". Curr Diab Rep 17, 95. 10.1007/s11892-017-0915-y.

45. Rahier, J., Wallon, J., and Henquin, J.C. (1981). Cell populations in the endocrine pancreas of human neonates and infants. Diabetologia 20, 540–546. 10.1007/BF00252762.

46. Saunders, D.C., Hart, N., Pan, F.C., Reihsmann, C.V., Hopkirk, A.L., Izmaylov, N., Mei, S., Sherrod, B.A., Davis, C., Duryea, J., et al. (2024). Mapping histological and functional maturation of human endocrine pancreas across early postnatal periods. bioRxiv. 10.1101/2024.12.20.629754.

47. Leete, P., Mallone, R., Richardson, S.J., Sosenko, J.M., Redondo, M.J., and Evans-Molina, C. (2018). The Effect of Age on the Progression and Severity of Type 1 Diabetes: Potential Effects on Disease Mechanisms. Curr Diab Rep 18, 115. 10.1007/s11892-018-1083-4.

48. Atkinson, M.A. (2014). Pancreatic biopsies in type 1 diabetes: revisiting the myth of Pandora’s box. Diabetologia 57, 656–659. 10.1007/s00125-013-3159-7.

49. Wasserfall, C., Montgomery, E., Yu, L., Michels, A., Gianani, R., Pugliese, A., Nierras, C., Kaddis, J.S., Schatz, D.A., Bonifacio, E., and Atkinson, M.A. (2016). Validation of a Rapid Type 1 Diabetes Autoantibody Screening Assay for Community Based Screening of Organ Donors to Identify Subjects at Increased Risk for the Disease. Clin Exp Immunol, Epub ahead of print. 10.1111/cei.12797.

50. Renier, N., Wu, Z., Simon, D.J., Yang, J., Ariel, P., and Tessier-Lavigne, M. (2014). iDISCO: a simple, rapid method to immunolabel large tissue samples for volume imaging. Cell 159, 896–910. 10.1016/j.cell.2014.10.010.

