## Supplemental Table 1 for "3D imaging of human pancreas suggests islet size and endocrine composition influence their loss in type 1 diabetes"

**Table S1. Donor information**

ND: no diabetes, GADA+: glutamic acid decarboxylase 65 autoantibody positive, T1D: type 1 diabetes, N/A: not applicable, F: female, M: male

| Donor Group | nPOD ID | RRID | T1D Duration (years) | Age (years) | Reported Sex | Reported Race/ Ethnicity |
|---|---|---|---|---|---|---|
| ND | 6588 | SAMN40555580 | N/A | 29 | F | Non-Hispanic White |
| | 6598 | SAMN40555585 | N/A | 17 | M | Hispanic/Latino |
| | 6601 | SAMN40555588 | N/A | 20 | M | Non-Hispanic White |
| | 6605 | SAMN40555592 | N/A | 23 | F | Non-Hispanic Black/African American |
| | 6607 | SAMN40555594 | N/A | 17 | F | Non-Hispanic White |
| | 6611 | SAMN44486497 | N/A | 14 | M | Non-Hispanic White |
| | 6612 | SAMN44486498 | N/A | 23 | M | Non-Hispanic White |
| | 6615 | SAMN44486501 | N/A | 14 | M | Non-Hispanic White |
| GADA+ | 6569 | SAMN38117299 | N/A | 20 | F | Hispanic/Latino |
| | 6573 | SAMN33284291 | N/A | 24 | F | Non-Hispanic White |
| | 6575 | SAMN33284293 | N/A | 23 | M | Non-Hispanic White |
| | 6582 | SAMN38117302 | N/A | 22 | M | Non-Hispanic White |
| | 6596 | SAMN40555583 | N/A | 18 | M | Non-Hispanic Black/African American |
| T1D | 6550 | SAMN25652261 | 0 | 25 | M | Non-Hispanic White |
| | 6566 | SAMN33284286 | 2 | 15 | M | Non-Hispanic White |
| | 6578 | SAMN33284295 | 0 | 11 | F | Non-Hispanic White |
| | 6579 | SAMN33284296 | 1 | 13 | F | Non-Hispanic White |
| | 6593 | SAMN38117312 | 1 | 26 | M | Non-Hispanic White |
| | 6604 | SAMN40555591 | 3 | 33 | F | Non-Hispanic Black/African American |
